# Surface-enhanced Raman spectroscopy on the membranes for antimicrobial resistance testing

**DOI:** 10.1101/2025.07.12.664554

**Authors:** Vladimir Mushenkov, Evgeny Andreev, Alexander Nechaev, Alexander Poddubikov, Vladimir Kukushkin, Elena Zavyalova

## Abstract

Antimicrobial resistance is one of the top global health threats; it is associated with millions of deaths per year. Traditional and most commonly used antibiotic susceptibility tests are based on detection of bacterial growth and its inhibition in the presence of an antimicrobial. These tests typically take over 1-2 days to perform, so empirical therapy schemes are often administered before the proper testing. Rapid tests for antimicrobial resistance are necessary to optimize the treatment of bacterial infection. A combination of MTT test with Raman spectroscopy to provide 1.5-hour long test antimicrobial susceptibility determination requiring 10^6^-10^8^ CFU/mL of bacteria. Here the first rapid antibiotic susceptibility test for trace amounts of bacteria is described. The bacterial titer can be decreased down to 10^2^ CFU/mL using surface-enhanced Raman spectroscopy (SERS) of the MTT-treated bacteria caught with the silver coated track-etched membranes allowing the antimicrobial testing of low-titer bacterial samples within 1.5 hour.

**Graphical Abstract:** 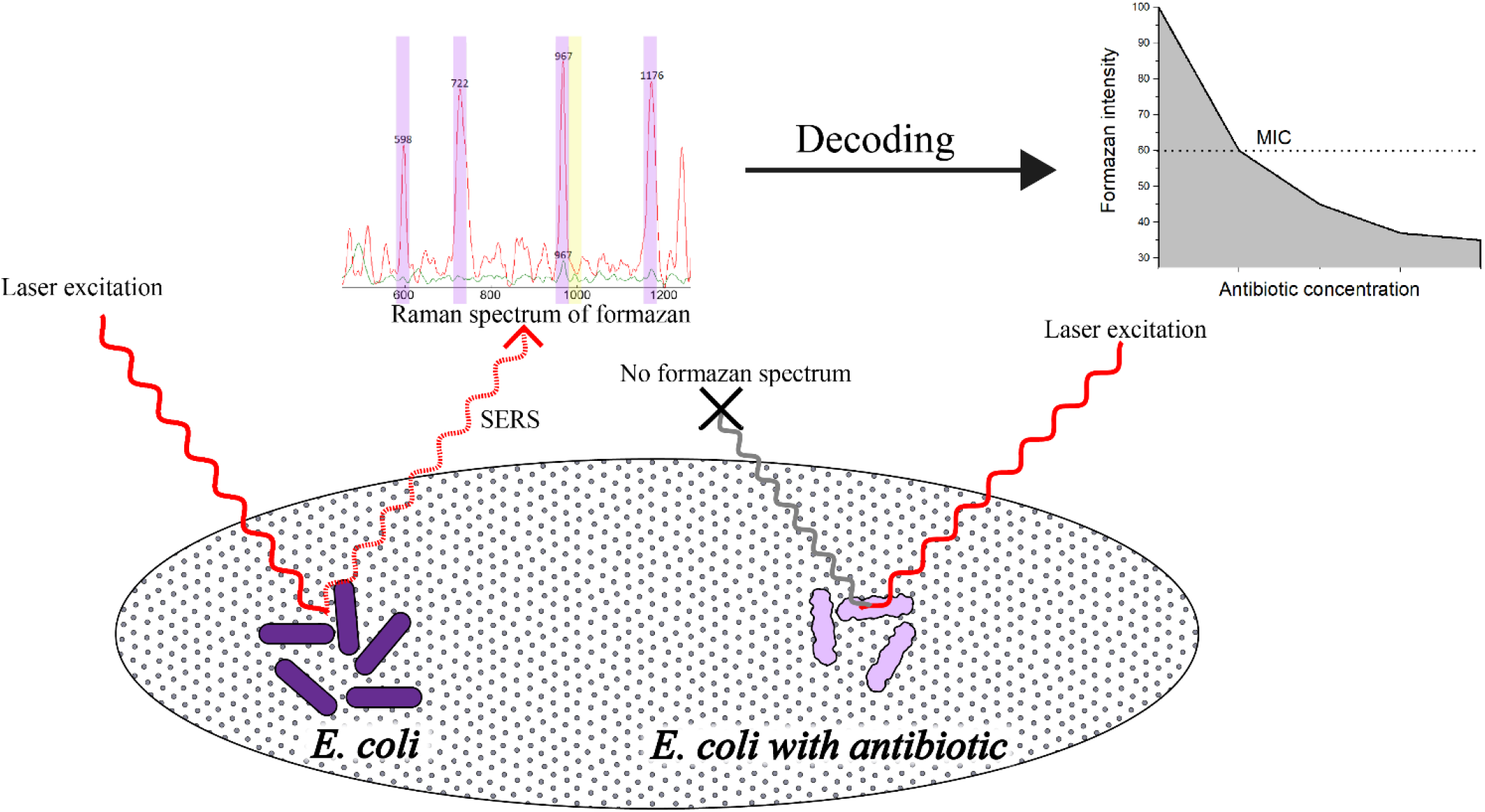

A combination of MTT test with membrane filtration and surface-enhanced Raman spectroscopy is used to determine
antibiotic susceptibility of *E.col*

## Introduction

Antibiotics, once introduced, revolutionized the treatment of infections, making many diseases much easier to cure. However, the effectiveness of antibacterial treatment is now continuously decreasing due to antibiotic resistance. Resistance is considered as one of the top global public health and development threats by World Health Organization [1]. At present times, bacterial antimicrobial resistance (AMR) is associated with nearly 5 million deaths annually, and mortality rates from drug-resistant infections could extent up to 80% in some cases [2,3]. Prompt administration of effective etiotropic therapy is essential for the treatment of drug-resistant infection [4]. Empirical therapy allows rapid applications, however in cases of resistant infections it shows significant failure rates 5, so antibiotic resistance tests (AST) are needed to select proper drug. In most cases, culture-based phenotypic ASTs are used, which are based on bacterial growth evaluation in the presence of the antibiotic, and takes over 1-2 days to perform [6,7]. These tests do not allow rapid resistance identification, so many alternative methods, which are based on different principles to achieve rapid resistance determination, are being developed [8].

MTT assay is a widely applied technique for assessing metabolic activity of the living cell. MTT assay is based on enzymatic reduction of tetrazolium salt (commonly 3-(4,5-dimethylthiazol-2-yl)-2,5-diphenyltetrazolium bromide, MTT) to reduced product formazan [9]. These enzymes are primarily NADH/NADPH-dependent dehydrogenases, whose activity is closely related to the cell metabolism level; thus, only active and viable cells will sustain efficient MTT reduction [10]. MTT assay could be used to determine antimicrobial activity [11] and detect resistant strains in much shorter time than cultural methods [12].

In MTT assay, absorbance spectrophotometry is commonly used for formazan quantification. However, it has some disadvantages due to an interference with the absorbance of the medium. Formazan extraction is necessary as an additional step [11]. Raman spectroscopy provides more reliable results due to registration of the unique spectrum, a “fingerprint” of the substance in complex media [13]. Raman spectrum of formazan reaches maximum intensity at excitation wavelength of 633 nm due to resonance Raman effect. Intensities of bands at 722 and 967 cm^-1^ have a linear dependence from formazan concentration and could be used to its quantification [14]. Previously, we developed a rapid antibiotic resistance test based on MTT assay and Raman spectroscopy (MTT-RS) [15]. This method evaluates minimal inhibitory concentration (MIC) in 1.5 h for a bacterial culture, providing reliable results which agrees well with classic cultural methods of resistance determination. However, the technique analyzes high concentrations of bacterial sample (2 McFarland units, approx. 6×10^8^ CFU/mL) requiring the preliminary culturing. The direct measurements of clinical cultures with characteristic bacterial amounts in the range of 10 - 1×10^7^ CFU/mL [16–18] are impossible. Therefore, increasing formazan detection sensitivity is needed for rapid point-of-care AST.

Surface-enhanced Raman spectroscopy (SERS) intensifies Raman spectrum due to surface plasmon resonance effect occurring when the analyte is in the proximity of rough metal surfaces (typically, silver or gold nanoparticles) [19]. When these surfaces are illuminated, excitation of collective electron oscillations, known as surface plasmons, emerges resonance which increases the intensity of Raman signal up to 10^10^-10^14^ times thus allowing single-molecule detection [20]. The simplest substrate for SERS is a colloid solution of nanoparticles, but their enhancement coefficients, in particular, due to aggregation on the biological matter [21].

Solid SERS substrates can provide more reliable and stronger signal due to larger hot spot quantity and complex structure providing stronger plasmon resonance [19]. As a solid substrate, permeable polymer membrane is a promising option as it can concentrate analyte and providing close connection to hot spots regardless of absorption or any other interactions [22]. Membrane methods are used in certified methods for determining pathogenic microorganisms as concentrators and work on the principles of a sieve separation mechanism based on the US Environmental Protection Agency recommendation (EPA -821-B-10-001; EPA-161; EPA-1622). Track membranes (TMs), obtained using ion-track technology, occupy a special place among membranes having uniformly sized, engineered nanopores formed via energetic heavy particle irradiation and chemical etching. Polyethylene terephthalate (PET) is commonly used due to its mechanical and chemical stability resulted in well-defined pore architecture and narrow pore size distribution that enable predictable transport and filtration properties [23,24]. TMs are widely used in medicine and biochemistry, in biological and chemical sensors, optical sensors for detection of enterobacteria and protozoa in water [25-29]. SERS technique could be coupled with TM speeding up pathogen identification [28,30,31]. Here we used this approach to increase sensitivity of AST.

## Materials and methods

Detailed information on reagents and methods is provided in the Supplementary materials.

## Results

Two types of PET membranes were studied. Membranes with cone-shaped pores were designed to catch the smallest microorganisms including mycoplasma (0.2-0.3 μ) and human viruses (0.1-0.3 μ). The membranes have the pore diameter of 570 nm at the side A and 110 nm at the side B (Figure 1). The low pore size dispersity allows creation of a filtration element with programable properties.

**Fig. 1.**
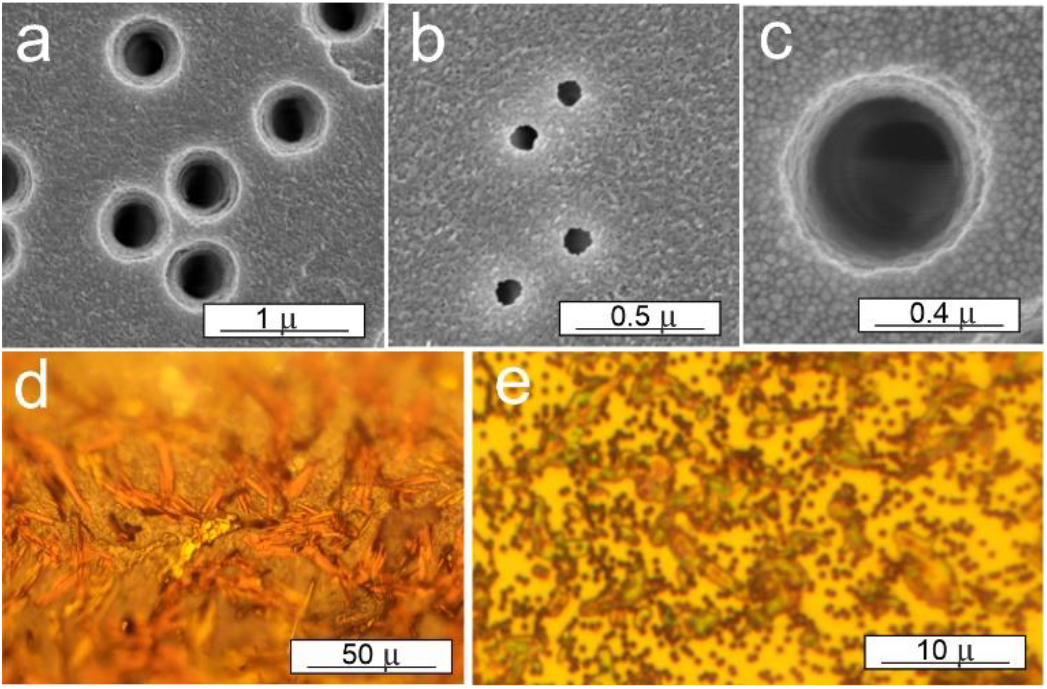
Microscopic images of track-etched membranes with cone-shaped pores. SEM images of the sides A (a) and B (b), the side A after thermal deposition of metals (c). Microscopic images of formazan crystals on the membrane (d) and separate *E.coli* cells stained with formazan (e).

Thermal deposition of Ag was carried out according to the previous study, where thin layers of Cr (10 Å) and Ag (80 Å) were thermally deposited on the membrane [27]. Cr increases adhesion of Ag particles to the polymer, whereas thin Ag layers formed nanoparticles that provided SERS-effect. The sample of *E.coli* K-12 strain after MTT-RS analysis (6×10^8^ CFU/mL) was filtered through this membrane providing a layer of formazan crystals (needle-like violet crystals, Figure 1f). The sample has characteristic formazan spectrum [15] which did not appear if metal-free membrane was used (Figure 2a). When the bacteria titer was decreased, the SERS intensity was also diminished. The crystals were not observed while the separate bacteria stained with formazan were clearly seen (Figure 1g). The spectra of the separate bacteria were acquired using 100x microscope lens (Figure 2b).

**Fig. 2.**
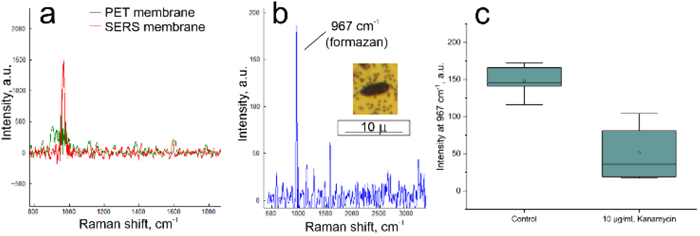
Formazan spectra on the SERS membrane with cone-shaped pores. (a) Comparison of spectra of MTT-treated *E.coli* K-12 strain cells obtained from PET and SERS membranes. (b) Spectrum of a single *E.coli* K-12 strain cell (shown in the inset) on the membrane obtained with 100x lens of SERS microscope. (c) The effect of the antibiotic on the formazan band intensity of separate *E.coli* cells.

The possibility of single cell study provides a unique opportunity to collect statistics of cell responses instead of an integral characteristic. Similar results can be obtained using flow cytometry; membrane-based technique is much simpler and excludes impurities in the system taking membranes and syringes as disposal elements. Figure 2c shows the first experiment on antibiotic effect on formazan spectrum intensity. Kanamycin-susceptible strain K-12 of *E.coli* in 5×10^7^ CFU/mL titer was treated with 10 μg/mL kanamycin for 60 min and then with MTT for another 30 min. The spectra of membranes with antibiotic-treated cells were about 3 times less intensive compared to the control untreated cells. This is the first in class data on SERS-membrane usage for AST.

Further experiments were conducted on another type of membrane. PET membrane with cylindric 0.2 μ pore (Figure 3) were used having good sterilization characteristics and apparatus-free filtration mode. This pore size allows retaining the majority of clinically-relevant bacteria types [32]. For example, two *E.coli* strains, M-17 and K-12 (approx. 8 10^3^ CFU/mL), were completely retained by the membrane (Figure 3). The membrane surface was coated with metals using the same protocol as for cone-shaped pore membranes. Next, a series of experiments were conducted to find a limit of detection (LoD) for this method and determine MIC for an antibiotic.

**Fig. 3.**
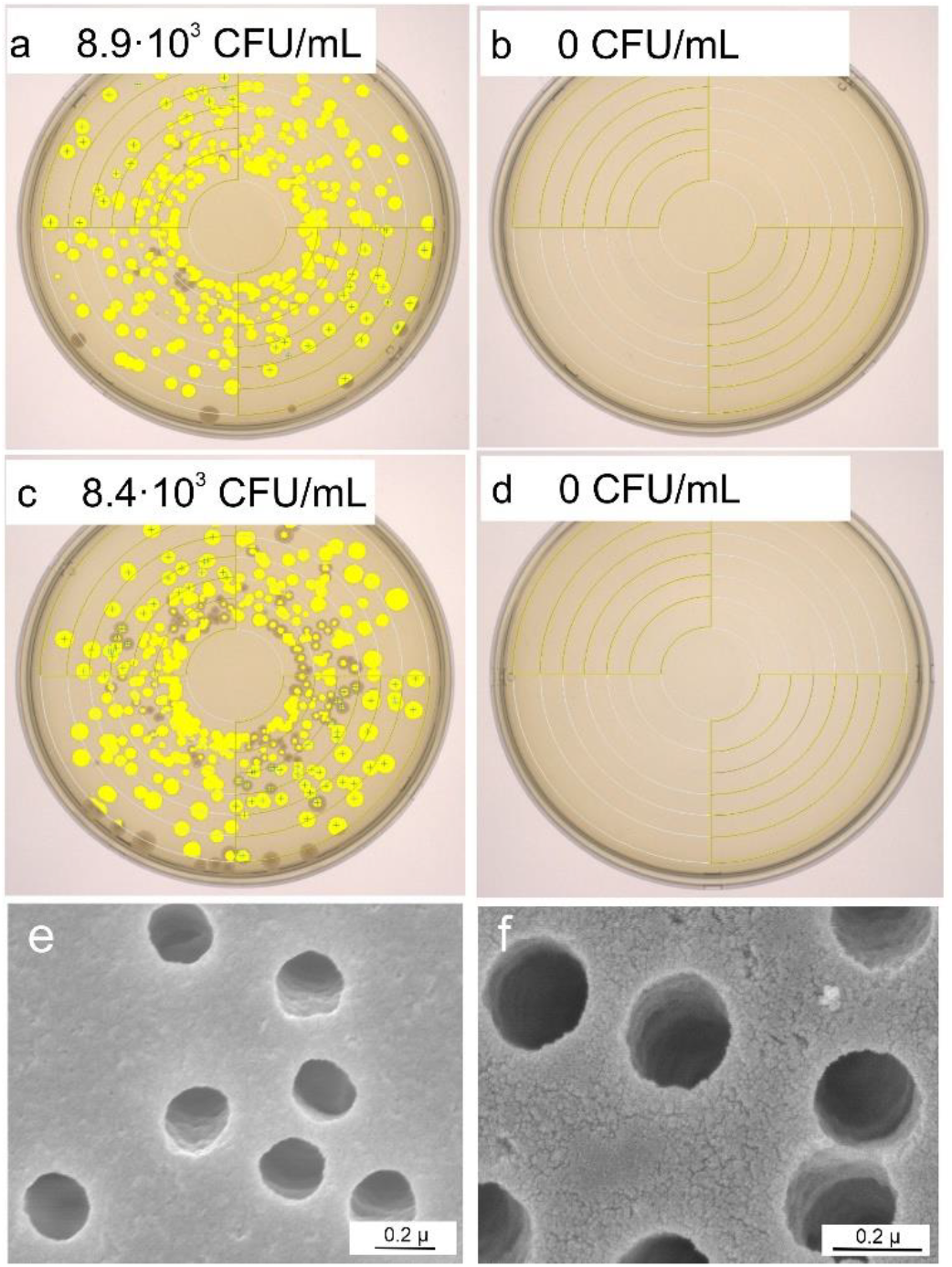
Efficiency of membrane filtration with 0.2 μ cylindric pore membranes. The number of colonies forming units were estimated for two *E.coli* strains, M-17 (a,b) and K-12 (c,d), before (a,c) and after (b,d) membrane filtration. SEM images of the membrane before (e) and after (f) thermal deposition of metals.

The new membranes allow obtaining of SERS spectra of the separate bacteria just as the first one (Figure 4). *E.coli* strain ATCC25922 was sequentially diluted to provide a series of samples with different bacteria amounts in the range of 10^2^-10^6^ CFU/ml. All dilutions were successfully detected having nearly the same formazan intensity for the single cell and a linear dependence of the spectra intensity on bacterial titer for large-area measurements (Figure 4). The LoD was estimated as 100 CFU/ml as further dilution provided sporadic cells in the microscope field with non-confident differences from the blank. Notably, the spectrum intensity of the single cell slightly increased with the decrease of bacterial titer. This result could be due to the increased number of MTT molecules per one cell. Large-area measurements were conducted further.

**Fig. 4.**
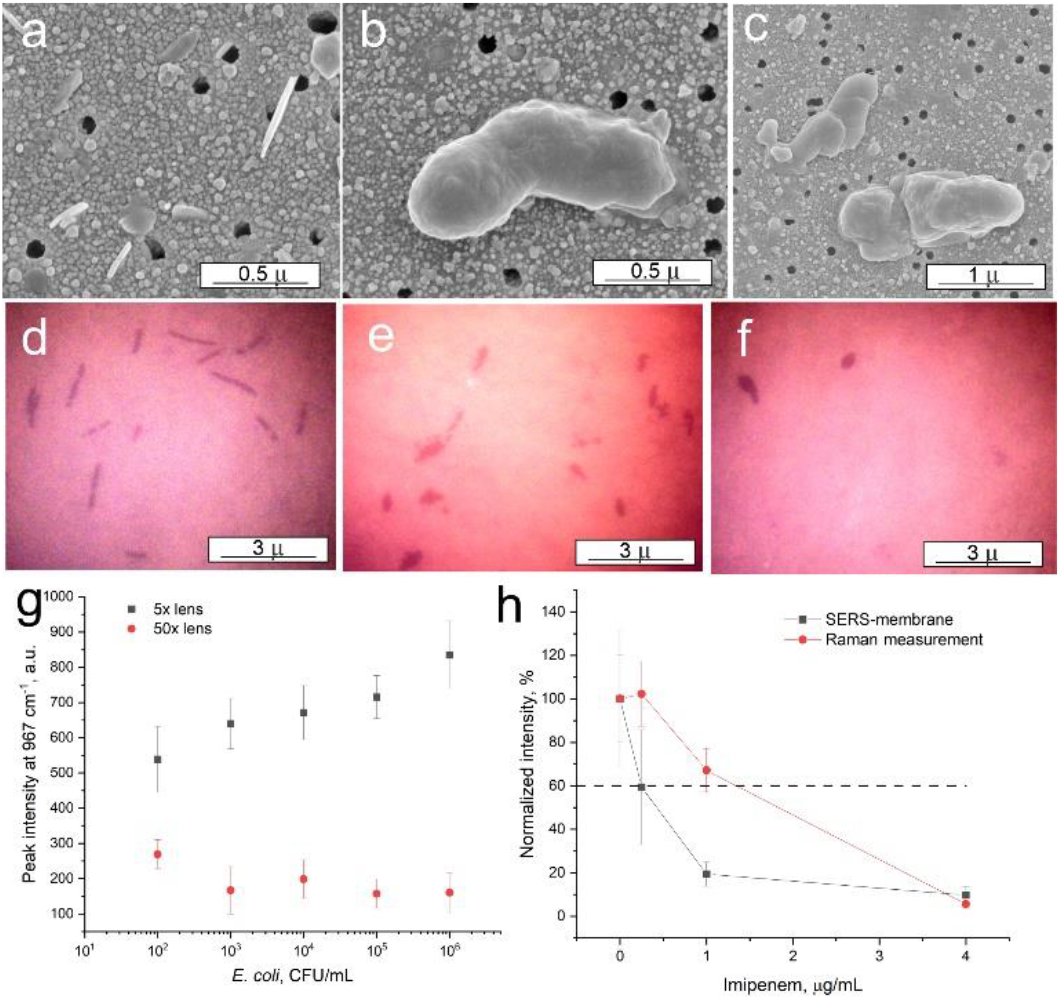
AST on SERS-active membrane with 0.2 μ cylindric pores. SEM images of formazan crystals (a), *E.coli* ATCC25922 strain cells (10^6^ CFU/mL) in the absence of antibiotics (b), *E.coli* ATCC25922 strain cells (10^6^ CFU/mL) in the presence of 32 μg/mL kanamycin. Microscopic images of *E.coli* ATCC25922 strain (5 10^5^ CFU/mL) after MTT treatment without antibiotic (d) either with 0.25 μg/mL (e) or 1 μg/mL (f) of imipenem. The limit of detection determination (g) and signal dependence on antibiotic concentration acquired for *E.coli* ATCC25922 strain (10^6^ CFU/mL) and imipenem (h) are shown.

Next, AST was conducted with *E.coli* ATCC25922 strain that is susceptible to imipenem and kanamycin. High concentrations of alive bacteria produced formazan crystals (Figure 4a). The cell morphology was retained after treatment with MTT (Figure 4b,d), whereas antibiotics disrupted cell morphology with no formazan staining of the cells (Figure 4c,e.f). The first experiment of MIC determination is shown in Figure 4h. MIC was estimated as antibiotic concentration caused 40%-drop in formazan spectrum intensity based on our previous experience in MTT-RS. Imipenem MIC determined with SERS-active membrane was 0.25 μg/mL, whereas the same value determined with MTT-RS was 1.25 μg/mL. The reference MIC from EUCAST is 0.125-0.25 μg/mL [38], so SERS-active membranes provided even more accurate result compared to MTT-RS.

Finally, *E.coli* ATCC25922 strain (50 CFU/mL) was incubated with 4 μg/mL of kanamycin to test the new AST on low-titer bacterial probes. The antibiotic decreased formazan band intensity to 70±20%. Thus, AMR of low-titer bacterial probes also can be studied, but the criteria should be elaborated further.

## Discussion

Broth microdilution, agar dilution, and Etest gradient strips are the standard methods for MIC determination. Dilution methods evaluate MIC by growing bacterial culture on media (liquid broth or agar plates) with increasing antibiotic concentrations, and MIC is considered as the first concentration at which the bacteria do not grow. These tests are the most accurate because they directly measure antibiotic resistance as the ability of bacteria to grow on media with antibiotic, and have minimal factors which can affect results, compared to other methods. The disadvantages of these methods are their labor intensity and test duration, typically ranging between 18-24h to perform the test for *E.coli* [34].

Etest is less labor-intensive and complicated method. An Etest plastic strip with antibiotic concentration gradient is placed on an agar plate with inoculated bacterial culture. MIC is then estimated as the concentration at which the inhibition zone disappears. Test duration is similar to dilution methods, but accuracy is lower in some cases [35]. Etest requires more concentrated bacterial culture (0.5-1 McFarland units, around 10^8^ CFU/mL, while dilution methods require 10^4^ – 10^5^ CFU/mL [34]).

Also, there are several automated systems for antimicrobial resistance testing which can estimate MICs, such as Vitek 2 or BD Phoenix. These systems greatly improve performance by automatizing most test steps and reducing test duration to 4-18h [36]. However, these systems are less reliable than dilution methods and Etest, and have a limited range of bacteria and antibiotics which can be tested [37]. These limitations, as well as the high cost of equipment and consumables, restrict these systems from widespread usage.

Most rapid and emerging AST methods determine only resistance without MIC data, but there are several methods based on microfluidics or light scattering, which can assess MIC in several hours [38]. A microfluidic test based on single cell growth rate measurements can provide MIC in 25 minutes with cell culture concentration of 10^4^ CFU/mL [39]. These rapid methods are not well-studied and their applicability, reliability and accuracy are not yet established. SERS-active membranes described here provide a unique possibility to study trace amounts of bacteria omitting cell culturing and making AST in point-of-care format.

## Conclusions

The first rapid antibiotic susceptibility test for trace amounts of bacteria is described. The bacterial titer can be decreased down to 10^2^ CFU/mL using surface-enhanced Raman spectroscopy (SERS) of the MTT-treated bacteria caught with the silver coated track-etched membranes allowing antimicrobial testing of low-titer bacterial samples within 1.5 hour.

## Acknowledgments

This work was supported by the Russian Science Foundation, grant no. 24-65-00015, https://rscf.ru/project/24-65-00015. The study was carried out on the scientific equipment of the Collective Usage Center “I.I.Mechnikov NIIVS”, Moscow, Russia. The authors thank O. Orelovich for the help with SEM examination of track-etched membrane samples.

