## Supplementary materials for "Surface-enhanced Raman spectroscopy on the membranes for antimicrobial resistance testing"

### Materials and methods

*Reagents*

MTT (3-(4,5-dimethylthiazol-2-yl)-2,5-diphenyltetrazolium bromide) was purchased from Dia-M (Russia). MTT was dissolved in water to final concentration of 4 g/L and stored at 4°C for a week. LB (Lennox broth) from Condalab (Italy), phosphate–buffered saline (PBS) with pH 7.3 from Ecoservice (Russia), kanamycin from Sigma–Aldrich (USA) and imipenem from Solarbio (China) were used.

*Track-etched membranes*

Two types of track-etched membranes were used in the work. Membranes with cylindrical pores and membranes with asymmetric cone-shaped pores. Track-etched membranes were manufactured at the Flerov Laboratory of Nuclear Reactions of the Joint Institute for Nuclear Research from Hostaphan brand PET by Mitsubishi Polyester Films (Japan) with a thickness of 19 and 23 µm. The membrane preparation included by irradiation with heavy ions at a cyclotron followed by sensitization with an ultraviolet radiation and chemical etching in sodium hydroxide solutions [1]. Membranes with cylindrical pores were etched in 2 M NaOH at 80°C [2]. An asymmetric track-etched membrane with cone-shaped pores was obtained using the method [3], the difference in obtaining such membranes was in irradiating the film with ultraviolet light on one side and etching the membrane in 5 M NaOH with the addition of 0.0125 wt.% Dowfax anionic surfactant 2A1 (Dow Chemical) at 60 °C. Characteristics of TMs used in this work are presented in Table 1.

Table 1. Characteristics of track-etched membranes.

| Pore shape | Pore diameter, µm | Membrane thickness, µm | Pore density, $cm^{-2}$ |
|---|---|---|---|
| Cylindrical | 0.20 | 23 | $4.5 \cdot 10^8$ |
| Cone-shaped | Side A 0.57 | 19 | $1 \cdot 10^8$ |
| | Side B 0.11 | | |

*Thermal sputtering of silver*

Sputtering was carried out on a NANO 38 thin film coating system (Kurt J. Lesker Company, Jefferson Hills, Pennsylvania, USA), a 10 Å chromium layer and a 80 Å silver layer were applied sequentially. Both layers were sprayed at a pressure of $8 \times 10^{-7}$ Torr with a deposition rate of 0.4 Å/c.

*Scanning electron microscopy*

The surface morphology and pore diameter of the membranes were studied by scanning electron microscopy (SEM) using a microscope SU8020 (Hitachi, Japan) with a cold field emission cathode. In order to improve the resolution and contrast of images, a thin layer of the Pt-Pd alloy was deposited on the samples.

*Bacteria*

Bacterial strains and culturing. Escherichia coli (*E.coli*) strains #03190 (M-17), #39030 (K-12); ATCC 25922 were obtained from the Collection of the «I.I.Mechnikov Center for Collective Use", Moscow, Russia. (USF Collection of the I.I.Mechnikov National Research Institute of Physics and Technology). The nutrient media were prepared and dispensed using the MEDIAWEL 10 automatic medium maker and the DISTRIWEL 440 dispensing module (France). The second passage of the daily culture was used. The working dilutions were prepared from a bacterial suspension of 0.5 McF, Densi-La-Meter II, Erba (Czechia). To accurately determine the number of viable cells in the bacterial suspension dilutions, direct seeding was performed followed by CFU counting. The bacterial suspension was seeded in a spiral seeding mode in a volume of 50 µL using the easySPIRAL Pro automatic seeding station, Intersciense (France), on Mueller Hinton Agar plates

(HiMedia, India). Bacterial cultures were grown for 16 hours at a temperature of 37°C in the Thermo B-20 incubator (Germany). The results (CFU/mL) were counted using the Scan4000 automatic colony counter, Intersciense. (France).

*SERS experiments*

Single colonies from each plate were cultured in LB overnight, diluted with another portion of LB to desired concentrations and incubated for 1h at 37°C. Bacterial concentration was calculated from the optical density of the culture at 600 nm wavelength. For MTT-RS test, samples of LB media (350 μL each) with different concentrations of antibiotic were prepared. In each sample we then added 50 μL of *E. coli* ATCC 25922 culture with 2 McFarland units of concentration. After 60 minutes of incubation at 37°C, we added 100 μL of MTT solution, stirred and then incubated for another 30 minutes before Raman spectroscopy.

Samples for SERS membrane testing were prepared similarly to MTT-RS experiment, with differences in *E. coli* strains (strains K-12 and ATCC 25922 were used) and titers, which were in the range of $50\text{-}5\cdot10^6$ CFU/mL. Each sample was then filtered with a syringe through a membrane locked in a filter holder. Filtration through cone-shaped pore membranes was performed using Elveflow Microfluidic device OB1 MK3+ with MFS flow sensor (Elvesys Microfluidics Innovation Center, France). Filtration trough 0.2 μ pore membranes was performed with a medical syringe manually. After sample filtration, 1 mL of PBS was filtered through the membrane to wash out residual MTT. For AST, the samples were prepared as for MTT-RS with lower bacterial titers.

For SERS measurements, RL637 and RL532 Raman spectrometers (Photon-Bio, Russia), integrated with PDV JX-40M microscope (Beijing PDV Instrument Co., Ltd, China) were used. Spectra were collected at 400 ms exposure time in 20 repeats. Raman spectra were collected with 10x and 100x lenses (for the 100x lens, the spectra acquisition area was focused on a single cell). For MTT-RS, we used RL637 Raman spectrometer without the microscope, on the same settings. Spectra were processed using Whittaker algorithm and asymmetrically reweighted penalized least squares smoothing [4,5].
